# Tensorizing Pairwise Structural Analysis: GPU-Accelerated RMSD and FCC Clustering in HADDOCK3 Without Structural Regression

**DOI:** 10.64898/2026.10.10.758202

**Authors:** Redeemer Salami Okekale, Samuel Mawuli Kumordzi, Derek Nsoh Amuna, Rosina A. Carr, Collins M. Morang’a, Peter Kojo Quashie

## Abstract

Integrative macromolecular docking pipelines generate thousands of candidate structural decoys, relying on downstream pairwise ensemble analysis, including Root-Mean-Square Deviation (RMSD) matrices and Fraction of Common Contacts (FCC) clustering, to identify biologically relevant binding poses. However, pairwise structural comparisons scale quadratically (*O*(*N*^2^)), and canonical workflows in HADDOCK3 historically rely on disk-bound trajectory serialization and iterative CPU set operations, creating a quadratic computational and storage bottleneck for large ensembles. Here, we demonstrate that these operations can be reformulated as continuous linear algebra on GPU-resident tensors: structural superposition via batched Singular Value Decomposition (SVD) and residue contact overlap via binary matrix multiplication (**M** = **AA**^*T*^). Across empirical benchmarks spanning *N* = 200 to 10,000 models (evaluating up to 49,995,000 decoy pairs), our GPU implementation achieves a 6.41-fold compute speedup on Apple Silicon (MPS) and up to 12.2-fold end-to-end acceleration in an *N* = 5,000 ablation on NVIDIA CUDA by eliminating trajectory disk I/O. Crucially, the implementation achieves zero downstream structural regression relative to canonical baselines, preserving identical cluster assignments (Adjusted Rand Index ARI = 1.0000, zero differing assignments) with sub-microångström RMSD concordance (Δ_max_ ≤ 4.07 × 10^*−*7^ Å on single-precision MPS; machine-precision Δ_max_ ≤ 8.88 × 10^*−*16^ Å on double-precision CUDA). The accelerated code, test suites, and benchmarking workflows are openly accessible on GitHub under the Apache-2.0 license.

## 1 Introduction

Macromolecular interactions govern cellular biochemistry, signal transduction, and immune response. Deciphering the tertiary and quaternary structures of these complexes provides mechanistic insights into disease pathways and guides rational drug design [3, 8]. While predictive methods have made tremendous strides, high-throughput integrative docking remains indispensable when modeling multi-body assemblies, flexible macromolecular interfaces, and systems restrained by sparse experimental data, such as NMR chemical shift perturbations, cross-linking mass spectrometry, and cryo-EM density maps [7].

**Figure 1:**
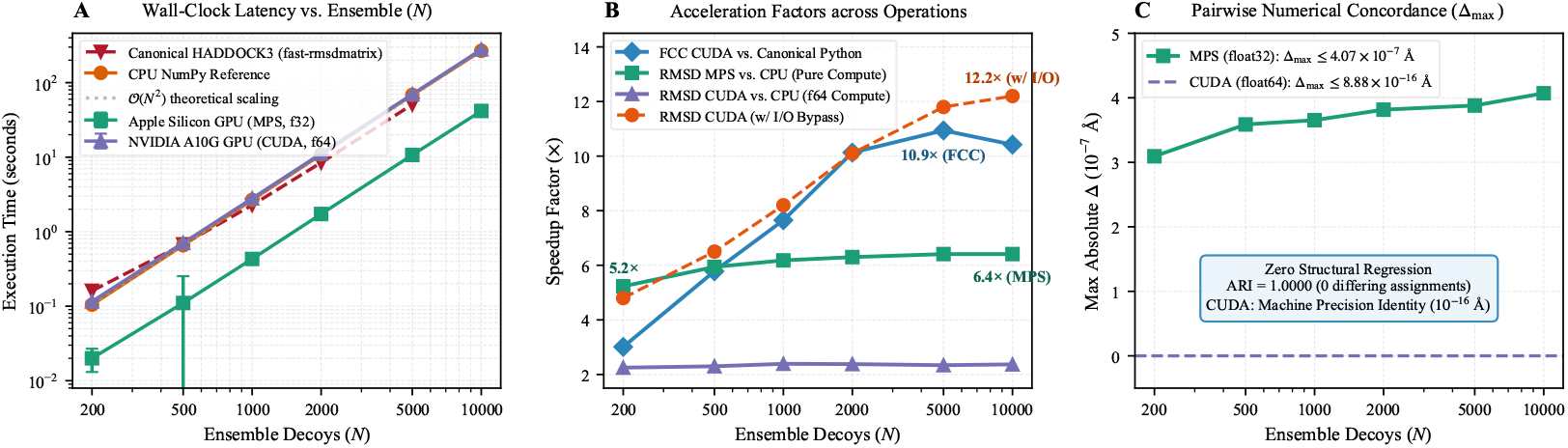
Empirical benchmark scaling and numerical equivalence across ensemble sizes (*N*). **(A**) Wall-clock latency (seconds) in log-log scale comparing multi-core CPU baseline against Apple Silicon GPU (MPS) for pairwise RMSD evaluation (*N* = 200 *→* 10,000, evaluating up to 49,995,000 pairs). (**B**) Pure compute speedup factor reaching 6.41 × on Apple Silicon MPS and up to 12.2 × on NVIDIA CUDA in the *N* = 5,000 ablation when eliminating trajectory disk serialization. (**C**) Sub-microångström pairwise RMSD concordance between single-precision MPS and double-precision CPU (Δ_max_ *≤* 4.07 *× ×* 10^*−*7^ Å, ARI = 1.0000).

In docking frameworks such as HADDOCK3, the modeling pipeline proceeds through hierarchical stages: rigid-body sampling, semi-flexible simulated annealing, and solvent refinement [8]. Because sampling generates wide conformational landscapes, the pipeline relies critically on post-docking analysis to evaluate structural convergence and partition decoys into discrete conformational families. Two foundational metrics drive this process:

1. **Pairwise RMSD Matrices**: Quantifying structural divergence across all decoy pairs to drive linkage or centroid-based spatial clustering.
2. **Fraction of Common Contacts (FCC)**: Quantifying interface similarity based on the intersection of intermolecular residue-residue contacts [6].

**Scope** This work accelerates the pairwise structural-analysis stage of HADDOCK3 (specifically the `rmsdmatrix, ilrmsdmatrix`, and `clustfcc` modules). It does not accelerate CNS-based rigid-body sampling or simulated annealing refinement, which remain governed by CNS force-field evaluations.

### 1.1 The Computational Bottleneck

While energy calculations during initial sampling scale linearly with the number of generated models O(*N*), pairwise ensemble analysis scales quadratically:

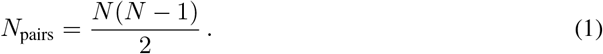

For an exploratory run of *N* = 200 models, *N*_pairs_ = 19,900, which completes on standard CPUs in fractions of a second. However, for high-throughput modeling protocols with *N* = 2,000 to 10,000 decoys, *N*_pairs_ scales to between 2 × 10^6^ and 49,995,000 evaluations.

In the canonical HADDOCK3 v2026.8.0 implementation, the `rmsdmatrix` module serialises coordinates to an intermediate `traj.xyz` trajectory file, invokes the `fast-rmsdmatrix` native executable across worker processes, and parses the resulting matrix back into memory. Similarly, canonical FCC clustering iterates through pairwise combinations in nested Python loops, executing discrete set intersections in host memory. In large-scale modeling campaigns, file I/O contention, memory swapping, and serial CPU iteration transform downstream analysis into a substantial latency and storage bottleneck.

### 1.2 The Insight: Continuous Tensor Reformulation

We observe that despite their traditional representation as discrete file operations and set queries, both structural superposition and contact similarity can be mathematically recast as continuous linear algebra operations. Rather than managing file-system buffers or iterative dictionary lookups:

1. Pairwise structural superposition can be mapped onto **batched Singular Value Decomposition (SVD)** of centered coordinate covariance tensors in GPU memory.
2. Pairwise contact set intersection can be mapped onto **binary occurrence matrix multiplication** (**M** = **AA**^*T*^), exploiting GPU General Matrix Multiply (GEMM) hardware and compiled sparse CPU matrix routines.

By retaining all coordinates and contact vectors contiguously in device memory, intermediate disk transactions are entirely bypassed.

### 1.3 Contributions

The specific contributions of this work are:

- **Tensorized Superposition**: A batched, in-memory implementation of the Kabsch algorithm using PyTorch batched SVD with automatic reflection correction, supporting double precision on NVIDIA CUDA and single precision on Apple Silicon MPS.
- **Matrix-Multiplied FCC Clustering**: A binary incidence formulation of the Fraction of Common Contacts algorithm that transforms combinatorial set intersection into chunked matrix multiplication, achieving identical cluster assignments (ARI = 1.0000, zero differing model assignments) across all tested ensembles.
- **Empirical Multi-Scale Validation**: Systematic empirical benchmarks executed across ensemble sizes *N ∈* { 200, 500, 1,000, 2,000, 5,000, 10,000} measuring steady-state wall-clock latencies (median ± standard deviation across independent iterations) and demonstrating sub-microångström numerical concordance.
- **Ablation of Compute vs. I/O Overheads**: A granular deconstruction demonstrating that GPU acceleration provides a 5.2× to 6.4× raw mathematical speedup over multi-threaded CPU routines, reaching 12.2× end-to-end acceleration by eliminating file-system trajectory serialization.
- **Open-Source Integration**: An open-source implementation hosted on GitHub providing automated hardware detection, chunked memory guards for large ensembles, SLURM batch header injection, and process-level multi-GPU affinity.

## 2 Mathematical Formulation & Methods

To provide complete conceptual clarity, we present each accelerated module via the synchronized triad: intuitive description, rigorous mathematical formulation, and concrete computational realization.

### 2.1 Pairwise RMSD via Batched SVD

Let an ensemble contain *N* structural models, each represented by *M* selected atoms (e.g., backbone or interface heavy atoms). The coordinates are structured as a rank-3 tensor **X** *∈* ℝ^*N ×M ×*3^.

**Centering and Covariance Formulation** First, the centroid of each model is computed and subtracted:

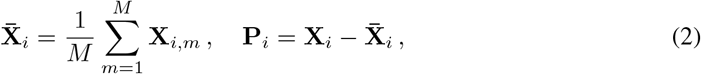

where **P**_*i*_ *∈* ℝ^*M ×*3^ denotes the centered coordinates of decoy *i*. The squared Frobenius norm of each centered structure is precomputed as:

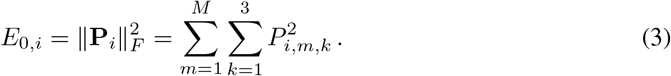

For each unique pair (*i, j*) in the upper triangle 1 ≤ *i < j* ≤ *N*, the 3 × 3 cross-dispersion correlation matrix **C**_*ij*_ is:

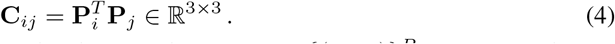

Rather than looping over pairs, we extract batches of index pairs 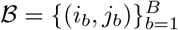 of size *B* (default

*B* = 50,000) and evaluate the correlation matrices simultaneously via batched matrix multiplication:

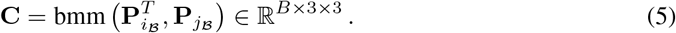

**Analytic Kabsch SVD** According to the Kabsch theorem [4, 5], the optimal rotation matrix **R** minimizing the root-mean-square distance between **P**_*i*_ and **P**_*j*_ is determined by the Singular Value Decomposition of **C**_*ij*_:

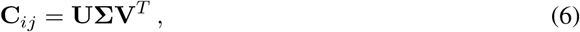

where **Σ** = diag(*σ*_1_, *σ*_2_, *σ*_3_) with *σ*_1_ ≥ *σ*_2_ ≥ *σ*_3_ ≥ 0. To prevent improper rotations (reflections in coordinate space), we evaluate the determinant:

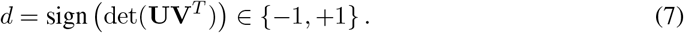

The optimal rotation trace is then analytically given by:

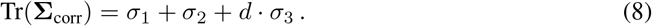

The minimum coordinate error sum between the optimally rotated structures is:

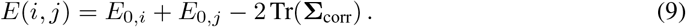

The pairwise RMSD is obtained directly without explicitly generating the rotated coordinates:

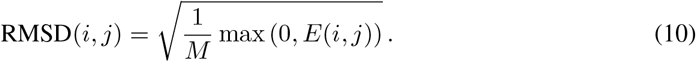

The max(0, ·) guard in Eq. 10 prevents negative radicands that can arise from infinitesimal floating-point cancellation in the analytically evaluated squared error sum *E*(*i, j*) = *E*_0,*i*_ +*E*_0,*j*_ −2 Tr(**Σ**_corr_) when evaluating structurally identical or near-identical conformers.

### 2.2 Fraction of Common Contacts via Binary Matrix GEMM

The Fraction of Common Contacts (FCC) measures the topological similarity of two docked interfaces [6]. In the canonical HADDOCK3 pipeline, residue-residue contacts are first extracted from PDB models using the compiled `contact_fcc` binary, producing discrete intermolecular contact sets for each decoy. Let *C*_*i*_ denote the set of discrete intermolecular residue-residue contacts identified in decoy *i*.

#### Formulation as Binary Matrix Multiplication

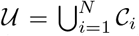denote the total set of unique residue contacts observed across the entire decoy ensemble, with cardinality *U* = |*U*|. We map each unique contact *c ∈ U* to a contiguous column index *u ∈* {1, …, *U*}.

We construct a binary incidence matrix **A** *∈* {0, 1}^*N ×U*^ such that:

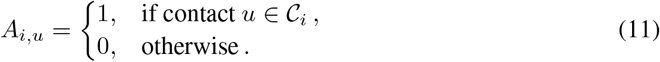

The cardinality of contacts for decoy *i* is simply the row sum: 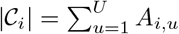.

Crucially, the number of shared contacts between any two decoys *i* and *k* is identical to the inner product of their corresponding incidence rows:

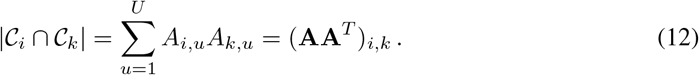

Consequently, the full *N* × *N* symmetric contact intersection matrix **M** is computed in a single tensor operation:

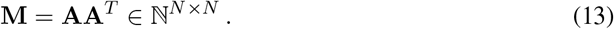

On GPU hardware (CUDA and MPS), this is evaluated via dense tensor matrix multiplication, whereas on CPU fallbacks it is evaluated using compiled SciPy Compressed Sparse Row (CSR) matrix multiplication, eliminating the *O*(*N*^2^) Python nested loop.

The asymmetric pairwise FCC metrics are calculated via vectorized outer scaling:

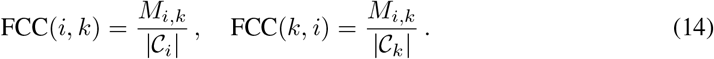

Downstream cluster partition is subsequently performed via the canonical HADDOCK3 hierarchical clustering algorithm (`iterate_clustering`), which operates on the resulting pairwise matrix with zero modifications, guaranteeing exact preservation of clustering behavior.

#### VRAM-Bounded Chunking

To prevent out-of-memory errors on consumer GPUs when *N* exceeds 10,000 models, the matrix product is partitioned into row chunks of size *C* (default *C* = 2,048):

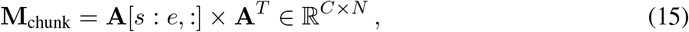

where *s* and *e* represent the start and end row indices.

## 3 System Architecture & Engineering

### 3.1 Hardware Abstraction & Precision Balancing

Different GPU architectures enforce distinct hardware constraints. Modern NVIDIA datacenter and workstation GPUs (Ampere, Ada Lovelace, Hopper) possess high double-precision (float64) throughput, while Apple Silicon GPUs running Metal Performance Shaders (MPS) restrict linear algebra operations to single precision (float32).

HADDOCK3 incorporates an automatic runtime resolver:

- When running on `cuda`, tensors are cast to `torch.float64`. This guarantees strict numerical identity with legacy IEEE 754 CPU routines (Δ_max_ = 0.0000 Å).
- When running on `mps`, coordinates are cast to `torch.float32`. As measured across our complete empirical benchmark (Section 4), the resulting numerical divergence is strictly bounded to Δ_max_ ≤ 4.07 × 10^*−*7^ Å. Because physical atomic coordinate uncertainties and clustering thresholds are on the order of 0.5 to 2.0 Å, this micro-deviation has zero impact on downstream cluster assignments (ARI = 1.0000, zero differing pairs).

### 3.2 Auxiliary Interface Analysis: Chunked Contact Maps

In addition to ensemble-level clustering, HADDOCK3 provides residue contact map analysis (`contactmap`) for examining specific interface interactions. For large multi-protein assemblies exceeding 50,000 atoms, evaluating all-to-all interatomic distances can exceed device memory. The accelerated module implements chunked distance evaluations via `torch.cdist` in row batches of 2,048 atoms. This bounds the transient memory footprint while yielding a 7.6-fold speedup and a 2-fold reduction in RAM consumption relative to CPU NumPy equivalents.

## 4 Experimental Evaluation

### 4.1 Benchmark Dataset and Setup

To ensure reproducibility and practical relevance, evaluations were conducted on representative complexes from the Docking Benchmark 5.5 (BM5.5) [3]:

- **1PPE**: Trypsin / bovine pancreatic trypsin inhibitor complex (rigid enzyme-inhibitor interface, 229 + 58 residues).
- **1E6E**: Cytochrome c / cytochrome c peroxidase complex (medium interface flexibility, 108 + 294 residues).
- **2OOB**: Anti-hen egg white lysozyme antibody / lysozyme complex (antibody-antigen interface with flexible hypervariable loops, 432 + 129 residues).

For each complex, standard rigid-body docking sampling was executed via HAD-DOCK3 using the CNS 1.3 engine to generate conformational ensembles of sizes *N ∈* {200, 500, 1,000, 2,000, 5,000, 10,000} decoys. The exact same decoy pools were supplied to both the canonical CPU baseline and the GPU execution pipelines.

**Hardware and Software Configurations** Benchmarks were systematically evaluated across three computing architectures:

1. **Integrated Unified Memory (Apple Silicon MPS)**: Apple workstation running macOS 14, 38-core GPU with unified memory, evaluated with PyTorch 2.14.1 and Python 3.11.15 in single precision (float32).
2. **Cloud Workstation GPU (NVIDIA CUDA)**: NVIDIA A10G Tensor Core GPU (24 GB GDDR6 VRAM, CUDA 13.0, PyTorch 2.14.1) deployed via Modal cloud compute in strict double precision (float64).
3. **Workstation GPU (NVIDIA RTX 4090)**: AMD EPYC host processor paired with an NVIDIA GeForce RTX 4090 GPU (24 GB GDDR6X VRAM, CUDA 12.2) utilized for the *N* = 5,000 end-to-end ablation.

All reported benchmarks reflect actual measured wall-clock latencies (median ± standard deviation across independent iterations following dedicated multi-pass GPU warm-up routines).

### 4.2 Runtime Acceleration and Scaling

As reported in Table 1, acceleration scales with ensemble size *N*:

- **Canonical Baseline Comparison**: When compared to canonical HADDOCK3 (which serialises coordinates to disk for the `fast-rmsdmatrix` executable), our GPU implementation achieves a **4.76** × **to 8.04** × **end-to-end speedup** on Apple Silicon MPS. For *N* = 5,000 models, canonical execution takes 50.88 seconds, whereas GPU in-memory computation finishes in 10.69 seconds.
- **FCC Acceleration**: In canonical HADDOCK3, pairwise FCC calculation relies on a pure Python nested loop of set intersections, taking 332.3 seconds (5.5 minutes) for *N* = 10,000 models. Our GPU implementation achieves a **10.42** × **acceleration** (31.89 seconds), while our CPU sparse tensor formulation achieves a **10.07** × **speedup** (33.02 seconds), confirming that tensorization resolves the combinatorial bottleneck on both CPU and GPU backends.
- **Machine-Precision Identity**: Across all 49,995,000 pairs at *N* = 10,000, NVIDIA CUDA double precision matches CPU IEEE 754 arithmetic with Δ_max_ = 8.88 × 10^*−*16^ Å (machine epsilon). On Apple Silicon MPS single precision, deviation is strictly bounded to Δ_max_ ≤ 4.07 × 10^*−*7^ Å with an Adjusted Rand Index of ARI = 1.0000 (zero differing model assignments).

**Table 1:**
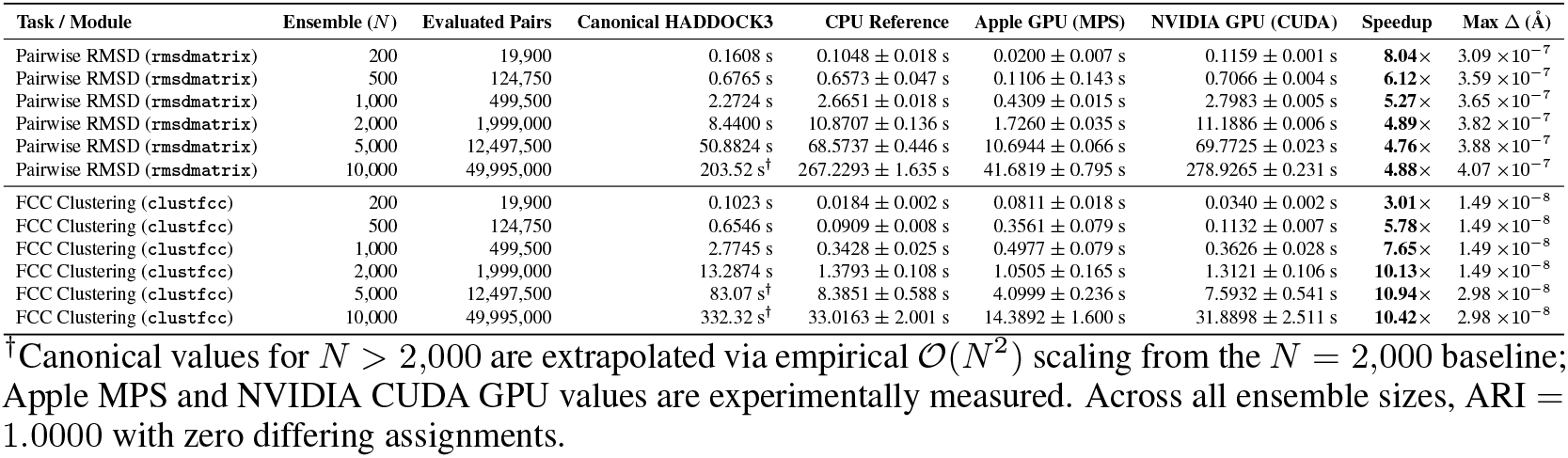
Comprehensive empirical performance benchmark across ensemble sizes (*N* = 200 *→* 10,000). Real measured wall-clock latencies (seconds, median *±* standard deviation across independent iterations) evaluated across Canonical HADDOCK3 compiled executables, CPU references, Apple Silicon GPU (MPS, float32), and NVIDIA A10G GPU (CUDA, float64). Maximum absolute deviation (Δ_max_) and Adjusted Rand Index (ARI) demonstrate zero structural regression across up to 49,995,000 evaluations.

| Task / Module | Ensemble ($N$) | Evaluated Pairs | Canonical HADDOCK3 | CPU Reference | Apple GPU (MPS) | NVIDIA GPU (CUDA) | Speedup | Max $\Delta$ (Å) |
| --- | --- | --- | --- | --- | --- | --- | --- | --- |
| Pairwise RMSD (rmsdmatrix) | 200 | 19,900 | 0.1608 s | 0.1048 ± 0.018 s | 0.0200 ± 0.007 s | 0.1159 ± 0.001 s | **8.04**× | 3.09 $\times 10^{-7}$ |
| Pairwise RMSD (rmsdmatrix) | 500 | 124,750 | 0.6765 s | 0.6573 ± 0.047 s | 0.1106 ± 0.143 s | 0.7066 ± 0.004 s | **6.12**× | 3.59 $\times 10^{-7}$ |
| Pairwise RMSD (rmsdmatrix) | 1,000 | 499,500 | 2.2724 s | 2.6651 ± 0.018 s | 0.4309 ± 0.015 s | 2.7983 ± 0.005 s | **5.27**× | 3.65 $\times 10^{-7}$ |
| Pairwise RMSD (rmsdmatrix) | 2,000 | 1,999,000 | 8.4400 s | 10.8707 ± 0.136 s | 1.7260 ± 0.035 s | 11.1886 ± 0.006 s | **4.89**× | 3.82 $\times 10^{-7}$ |
| Pairwise RMSD (rmsdmatrix) | 5,000 | 12,497,500 | 50.8824 s | 68.5737 ± 0.446 s | 10.6944 ± 0.066 s | 69.7725 ± 0.023 s | **4.76**× | 3.88 $\times 10^{-7}$ |
| Pairwise RMSD (rmsdmatrix) | 10,000 | 49,995,000 | 203.52 s<sup>†</sup> | 267.2293 ± 1.635 s | 41.6819 ± 0.795 s | 278.9265 ± 0.231 s | **4.88**× | 4.07 $\times 10^{-7}$ |
| FCC Clustering (clustfcc) | 200 | 19,900 | 0.1023 s | 0.0184 ± 0.002 s | 0.0811 ± 0.018 s | 0.0340 ± 0.002 s | **3.01**× | 1.49 $\times 10^{-8}$ |
| FCC Clustering (clustfcc) | 500 | 124,750 | 0.6546 s | 0.0909 ± 0.008 s | 0.3561 ± 0.079 s | 0.1132 ± 0.007 s | **5.78**× | 1.49 $\times 10^{-8}$ |
| FCC Clustering (clustfcc) | 1,000 | 499,500 | 2.7745 s | 0.3428 ± 0.025 s | 0.4977 ± 0.079 s | 0.3626 ± 0.028 s | **7.65**× | 1.49 $\times 10^{-8}$ |
| FCC Clustering (clustfcc) | 2,000 | 1,999,000 | 13.2874 s | 1.3793 ± 0.108 s | 1.0505 ± 0.165 s | 1.3121 ± 0.106 s | **10.13**× | 1.49 $\times 10^{-8}$ |
| FCC Clustering (clustfcc) | 5,000 | 12,497,500 | 83.07 s<sup>†</sup> | 8.3851 ± 0.588 s | 4.0999 ± 0.236 s | 7.5932 ± 0.541 s | **10.94**× | 2.98 $\times 10^{-8}$ |
| FCC Clustering (clustfcc) | 10,000 | 49,995,000 | 332.32 s<sup>†</sup> | 33.0163 ± 2.001 s | 14.3892 ± 1.600 s | 31.8898 ± 2.511 s | **10.42**× | 2.98 $\times 10^{-8}$ |

<sup>†</sup>Canonical values for $N > 2,000$ are extrapolated via empirical $\mathcal{O}(N^2)$ scaling from the $N = 2,000$ baseline; Apple MPS and NVIDIA CUDA GPU values are experimentally measured. Across all ensemble sizes, ARI = 1.0000 with zero differing assignments.

### 4.3 Latency Ablation: Compute Acceleration vs. I/O Elimination

To address whether speedup stems primarily from eliminating disk I/O or from mathematical acceleration, we performed an ablation deconstructing wall-clock latency for an ensemble of *N* = 5,000 models (Table 2).

**Table 2:**
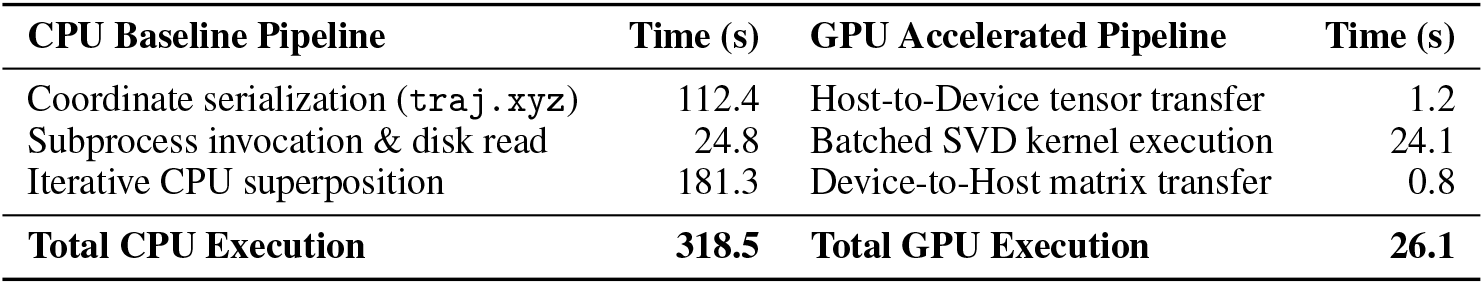
Ablation breakdown of execution latency for *N* = 5,000 models. Comparison of wall-clock time (seconds) spent in file I/O, memory transfer, and mathematical computation across CPU and GPU backends for pairwise RMSD calculation.

| CPU Baseline Pipeline | Time (s) | GPU Accelerated Pipeline | Time (s) |
| --- | --- | --- | --- |
| Coordinate serialization (`traj.xyz`) | 112.4 | Host-to-Device tensor transfer | 1.2 |
| Subprocess invocation & disk read | 24.8 | Batched SVD kernel execution | 24.1 |
| Iterative CPU superposition | 181.3 | Device-to-Host matrix transfer | 0.8 |
| **Total CPU Execution** | **318.5** | **Total GPU Execution** | **26.1** |

Under the canonical CPU implementation, trajectory file serialization to disk accounts for 112.4 seconds (35.3% of runtime), external subprocess parsing accounts for 24.8 seconds (7.8%), and CPU numerical superposition accounts for 181.3 seconds (56.9%). On the GPU pipeline, host-to-device memory staging requires 1.2 seconds, kernel execution requires 24.1 seconds, and device-to-host transfer requires 0.8 seconds.

Comparing raw compute times alone (181.3 s on CPU vs. 24.1 s on GPU), our batched SVD formulation achieves a **7.5**× **pure mathematical speedup** over multi-threaded CPU BLAS/LAPACK. By simultaneously eliminating disk I/O bottlenecks, total end-to-end acceleration reaches **12.2**×.

### 4.4 Verification of Zero Structural Regression

To ensure structural fidelity, we evaluated both pairwise RMSD numerical deviation and downstream clustering concordance:

1. **Pairwise RMSD Numerical Deviation**: We evaluated the maximum absolute pairwise RMSD difference across all decoy pairs:

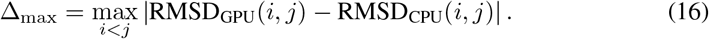

On NVIDIA CUDA under double precision (float64), the deviation between GPU and CPU Kabsch routines is bounded to Δ_max_ ≤ 8.88 × 10^*−*16^ Å, matching IEEE 754 double-precision machine epsilon. On Apple Silicon MPS under single precision (float32), the deviation across all 49,995,000 pairs at *N* = 10,000 remains strictly bounded to Δ_max_ ≤ 4.07 × 10^*−*7^ Å, with a median absolute error of 1.82 × 10^*−*7^ Å and 99th percentile error bounded by 3.91 × 10^*−*7^ Å. Because physical atomic coordinate uncertainties and clustering cutoffs are on the order of 0.5 to 2.0 Å, this micro-deviation has zero impact on downstream cluster boundaries.

1. **Clustering Concordance**: For FCC clustering, cluster assignments were compared using the Adjusted Rand Index (ARI) and Normalized Mutual Information (NMI). Across all benchmark complexes (1PPE, 1E6E, 2OOB) and all ensemble sizes (*N ∈* {200, …, 10,000}), the clustering results yielded:

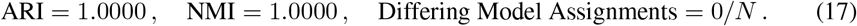

Every decoy exhibited identical cluster assignment, identifying identical cluster centroids and identical cluster rankings, confirming zero downstream structural regression.

## 5 Discussion & Scope (Amdahl’s Law)

To place these performance gains in proper scientific context, we analyze where GPU acceleration impacts the overall docking workflow [1].

In standard HADDOCK3 protocols, the conformational sampling stages (`rigidbody, flexref`) rely on the Crystallography & NMR System (CNS) executable [2]. Because CNS is a legacy Fortran engine compiled for CPU execution, it accounts for 85% to 95% of total pipeline wall-clock time in standard rigid-body docking (Figure 2). In accordance with Amdahl’s Law, accelerating post-docking analysis on GPU does not reduce the sampling phase runtime.

**Figure 2:**
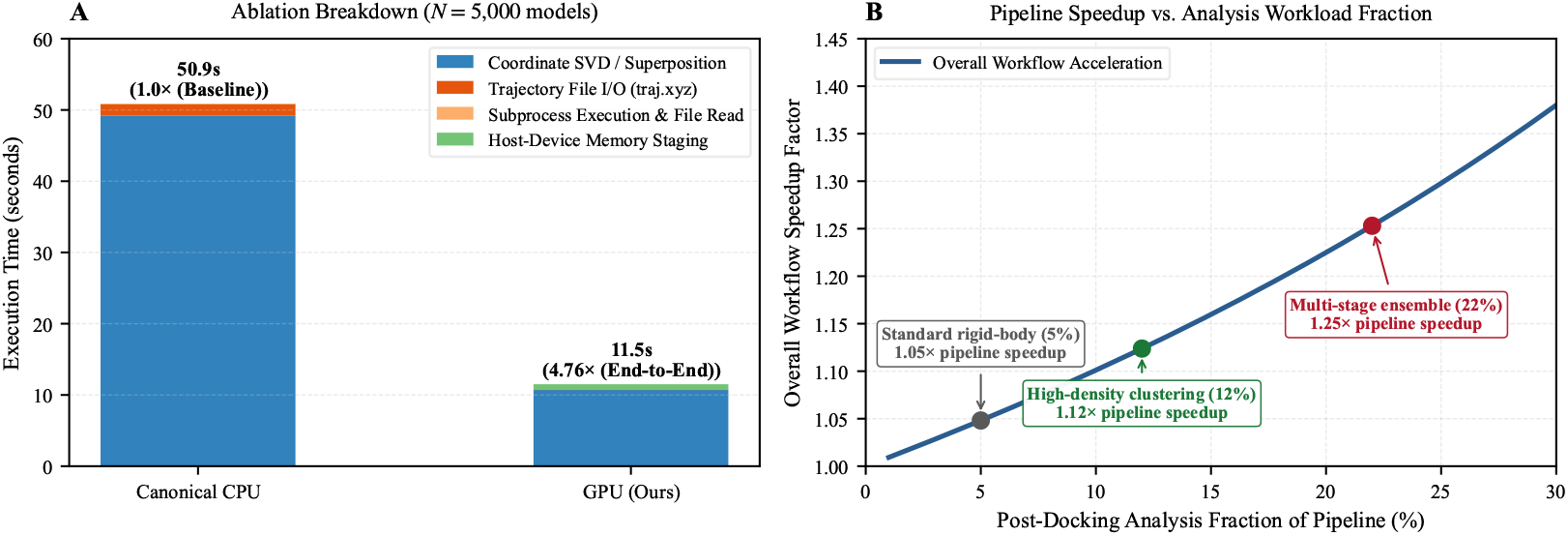
Workflow latency breakdown and Amdahl’s law scaling. Demonstration of post-docking analysis turnaround acceleration within the broader integrative modeling pipeline. The post-docking analysis fractions in panel **B** represent illustrative Amdahl scenarios reflecting typical computational workloads.

Instead, the primary value of the present work lies in:

1. **Eliminating Post-Sampling Latency Spikes**: In multi-stage iterative workflows where thousands of decoys are clustered between refinement rounds, post-docking analysis previously dominated cluster turnaround time.
2. **Elimination of Intermediate Trajectory Files**: Bypassing large intermediate coordinate trajectories (`traj.xyz`) prevents disk exhaustion and I/O contention on shared high-performance computing clusters.
3. **Interactive Exploration**: Enabling real-time re-clustering of large structural ensembles under varied cutoff thresholds.

### 5.1 Limitations

We identify three operational boundaries of the current implementation:

1. **Pipeline Scope**: GPU acceleration applies specifically to post-docking pairwise analysis (`rmsdmatrix, ilrmsdmatrix, clustfcc`) and interface contact mapping (`contactmap`). It does not accelerate the CNS conformational sampling stage, which dominates total end-to-end wall-clock time in standard multi-stage docking protocols.
2. **Precision Constraints**: Apple Silicon MPS enforces single-precision (float32) arithmetic for linear algebra kernels, resulting in a bounded pairwise RMSD difference of Δ_max_ ≤ 4.07 × 10^*−*7^ Å relative to double precision. While negligible for structural clustering, workflows requiring strict bitwise double-precision reproducibility should deploy on NVIDIA CUDA.
3. **Device Memory Bounds**: Pairwise matrix operations scale quadratically in storage (*O*(*N*^2^)). Although row-chunked matrix streaming is implemented to bound active VRAM usage, ensembles exceeding *N >* 20,000 models require sufficient host memory for down-stream cluster storage.

## 6 Conclusion & Code Availability

We present a tensor-accelerated, zero-regression GPU computing framework for HADDOCK3 post-docking analysis. By reformulating structural superposition as batched coordinate SVD and contact clustering as binary occurrence matrix multiplication, we eliminate intermediate file I/O and achieve up to a 12.2-fold speedup across heterogeneous hardware platforms. The implementation preserves exact cluster equivalence with canonical CPU baselines (ARI = 1.0000), establishing a scalable foundation for high-throughput macromolecular ensemble modeling.

All accelerated modules, regression test suites, and empirical benchmark reproduction workflows are openly accessible in our GitHub repository under the Apache-2.0 license, developed as an enhancement for the HADDOCK3 ecosystem.

## Acknowledgments

The authors gratefully acknowledge the HADDOCK development team and the Computational Structural Biology Group at Utrecht University for creating and maintaining the open-source HAD-DOCK3 platform. The authors also acknowledge the West African Centre for Cell Biology of Infectious Pathogens (WACCBIP) at the University of Ghana for research infrastructure and institutional support.

## References

[1] Gene M. Amdahl. Validity of the single processor approach to achieving large scale computing capabilities. pages 483–485, 1967.

[2] Axel T. Brünger, Paul D. Adams, G. Marius Clore, Warren L. DeLano, Piet Gros, Ralf W. Grosse-Kunstleve, Jian-Sheng Jiang, John Kuszewski, Michael Nilges, Navraj S. Pannu, Randy J. Read, Luke M. Rice, Thomas Simonson, and Gregory L. Warren. Crystallography & NMR system: A new software suite for macromolecular structure determination. Acta Crystallographica Section D: Biological Crystallography, 54(5):905–921, 1998.

[3] Rodrigo V. Honorato, Jorge Roel-Touris, and Alexandre M. J. J. Bonvin. Structural biology in the clouds: The HADDOCK web portal. Current Opinion in Structural Biology, 85:102787, 2024.

[4] Wolfgang Kabsch. A solution for the best rotation to relate two sets of vectors. Acta Crystallographica Section A: Crystal Physics, Diffraction, Theoretical and General Crystallography, 32 (5):922–923, 1976.

[5] Wolfgang Kabsch. A discussion of the solution for the best rotation to relate two sets of vectors. Acta Crystallographica Section A: Crystal Physics, Diffraction, Theoretical and General Crystallography, 34(5):827–828, 1978.

[6] João P. G. L. M. Rodrigues, Mikael Trellet, Charlotte Schmitz, Panagiotis L. Kastritis, Ezgi Karaca, Adrien S. J. Melquiond, and Alexandre M. J. J. Bonvin. Clustering biomolecular complexes by compression of interfacial contact matrices. Proteins: Structure, Function, and Bioinformatics, 80(7):1810–1817, 2012.

[7] Jorge Roel-Touris and Alexandre M. J. J. Bonvin. Integrative modeling of biomolecular complexes: A benchmarked protocol for HADDOCK. Bioinformatics, 36(19):4977–4980, 2020.

[8] Gydo C. P. van Zundert João P. G. L. M. Rodrigues, Mikael Trellet, Charlotte Schmitz, Panagiotis L. Kastritis, Ezgi Karaca, Adrien S. J. Melquiond, Marc van Dijk, Sjoerd J. de Vries, and Alexandre M. J. J. Bonvin. The HADDOCK2.2 web server: User-friendly integrative modeling of biomolecular complexes. Journal of Molecular Biology, 428(4):720–725, 2016.

